# Oral Silver Nanoparticle Exposure Increases Phosphorylated Tau Protein and Antioxidant Expression in Dentate Gyrus of Rat Brain

**DOI:** 10.64898/2026.11.23.768619

**Authors:** Rianna N. Smith, Eden S. Cantrell, William B. Crockett, Nadja Spitzer

## Abstract

Silver nanoparticles (AgNPs) are incorporated into products including toys, cosmetics, and surgical tools due to their antimicrobial properties. AgNPs can cross the blood-brain barrier and bioaccumulate in the brain, persisting for months with no clearance. We previously found that AgNPs disrupt the f-actin cytoskeleton, induce neurite collapse, and disrupt cytoskeletal regulatory pathways in cultured neural cells. The cytoskeleton is critical to maintaining synaptic structure and mediating neuroplasticity, therefore we investigated if AgNPs induce nuclear accumulation of hyperphosphorylated tau (ptau) and superoxide dismutase-1 (SOD1) in the dentate gyrus of the hippocampus to further assess nano-bio interactions in the brain. Following nine weekdays of treatment with 0, 25, or 100 mg/kg oral AgNPs, brains were collected one day after treatment to measure acute effects or one month post-treatment to measure the effect of bioaccumulated AgNPs. Using immunohistochemistry, we determined that ptau aggregates and SOD1 expression were significantly increased in the dentate gyrus after acute exposure and remained significantly elevated after exposure to bioaccumulated AgNPs. These findings suggest that exposure to AgNPs induces persistent disruptions to cytoskeletal regulatory proteins and antioxidant mechanisms associated with neurogenerative disease. Acute AgNPs exposure also induced minor disruptions in neurogenesis in the subventricular zone – rostral migratory stream – olfactory bulb niche that was reversed in rats exposed to bioaccumulated AgNPs. As AgNPs continue to be incorporated into everyday products, these data emphasize the importance of investigating their effect on mechanisms that may increase the risk of neurodegenerative diseases.

## INTRODUCTION

Silver nanoparticles (AgNPs) are widely incorporated in a variety of products including toys, cosmetics, and surgical tools due to their unique antimicrobial properties.^1–3^ Emerging evidence has spotlighted the potential negative effects of AgNPs release and persistence in the environment. Silver is released into environmental water systems from manufacturing facilities, ^4, 5^ and shed from products during use and after disposal. For example, toothbrushes release up to 10.2 ng of AgNPs per use,^1^ 87% of embedded silver, including AgNPs and ionic silver, is released from antimicrobial textiles after five washes,^3^ and wound dressings release 17-111 mg of silver over the first 48 hours after application.^3^ This leads to contamination of wastewater, surface water, and agricultural products,^1, 4, 6, 7^ and the eventual ingestion and chronic exposure to AgNPs in people and animals.^8–15^ The concentration of AgNPs in aquatic ecosystems has been estimated in the ng/L range.^16–18^ Environmental AgNP contamination persists over time,^15, 19^ and AgNPs bioaccumulate in the tissues of animals, moving through trophic levels in and around contaminated sites.^6, 15, 20^ In rodents, ingested AgNPs readily enter the body, disperse via the bloodstream, and accumulate in various tissues including brain.^21–23^ Moreover, accumulated silver persists in brain for months after oral AgNP exposure with very little clearance.^21, 22^

Here, we endeavored to model exposure representative of chronic environmental consumer exposure to AgNP-enhanced products, water, and via the food chain and agriculture.^15, 24, 25^ We used a nine-day dosing paradigm to achieve cumulative brain Ag levels in the ng/g tissue range. This represents a time-compressed approach to reach brain Ag levels thought to approximate the range predicted from repeated chronic exposure to environmental AgNPs combined with their, bioaccumulation, and persistence in neural tissue.^26^ Our treatments were also comparable to other rodent studies reporting ng/g tissue levels after repeated oral exposure.^27–29^ Our previous work showed that cultured mammalian cells did not suffer DNA fragmentation or cell death at doses below 1 μg/ml, but this dose did disrupt cellular organization, intracellular signaling pathways, and neurite structure.^30–33^ Similarly, the exposure model used here in rats did not induce overt signs of toxicity. We assessed rat brain tissue collected immediately after the acute treatment and tissue collected one month after treatment to determine the effects of exposure to bioaccumulated AgNPs on markers of neurodegeneration and on neurogenesis.

Many water filters cannot remove AgNPs, indeed, some water filtration systems include AgNPs to prevent microbial growth. Furthermore, the remarkable surface area and physicochemical properties of AgNPs allow them to easily pass biological barriers via endocytosis, transcytosis, paracellular transport, or passive diffusion.^34^ AgNPs enter the bloodstream from the gut after oral exposure^5^ and readily cross the blood-brain barrier (BBB),^5, 21, 22, 28, 35^ the brain’s first line of defense against toxins. They interact with the tight junctions that regulate and maintain the BBB, ultimately increasing its permeability and facilitating further entry of AgNPs into brain tissue.^34^ Once inside the brain, AgNPs bioaccumulate with no clearance for up to several months in mammals.^5^

Whereas tissues like liver can potentially compensate for damage from AgNP accumulation,^10^ the specificity of brain tissue and synaptic organization may be more susceptible to localized disruption by AgNP incorporation and persistence. AgNPs induce the generation of reactive oxygen species (ROS) in the brain, leading to oxidative stress.^36^ Oxidative stress is an established driving force behind the misfolding of proteins like tubulin associated unit (tau), a critical microtubule scaffolding protein that organizes tubulin and maintains cytoskeletal structure.^37^ Oxidative imbalance can induce hyperphosphorylation of tau (ptau), triggering a change in protein conformation and formation of harmful aggregates that cause cells to lose signaling capabilities and eventually apoptose.^38^ Oxidative stress can induce ptau, and accumulations of ptau can induce the generation of ROS in a positive feedback cycle.^39^ Tau aggregates spread throughout the brain via prion-like mechanisms,^40^ interacting with normal tau protein to induce disassociation from the microtubule and translocation to the nucleus.^40^ This destabilizes the cytoskeleton and promotes neurodegeneration^40^ associated with Alzheimer’s disease and other neurogenerative disorders.^41^ AgNPs have also been linked to other neurodegenerative mechanisms, promoting amyloid expression and formation of β-amyloid plaques in neural cells.^42^

Toxic ROS are broken down and converted into molecular oxygen and hydrogen peroxide by Copper/Zinc superoxide dismutase 1 (SOD1).^43^ Cytoplasmic SOD1 shifts to the nucleus during times of cellular stress to protect DNA.^44^ Elevations of ROS and SOD1 are precursors to neurodegenerative pathologies.^43^ In the brain, the hippocampal dentate gyrus (DG), which aids in processing associative memory, is altered during neurodegenerative disease.^45^ AgNP-mediated molecular disruptions within the DG may therefore have consequences extending beyond cellular stress and could influence the mechanisms underlying learning and memory. Thus, we used immunohistochemistry (IHC) to determine if exposure to AgNPs induces the hyperphosphorylation and nuclear translocation of tau protein and increases the expression of SOD1 in DG nuclei of rat brain.

Because microtubule cytoskeleton dynamics and stability are critical to cell division and migration,^40, 46^ we also assessed neurogenesis in the subventricular zone – rostral migratory stream – olfactory bulb (SVZ-RMS-OB) niche in rat brain after exposure to acute or bioaccumulated AgNPs. In mammals, the SVZ is one of two well characterized regions of adult neurogenesis.^47^ Neural cells born in the SVZ migrate along the RMS, continuing to proliferate until they reach the OB where they differentiate into granular and periglomerular interneurons.^48–50^ Incorporation of new neurons into OB pre-existing circuitry is essential for learning and recognizing odorants.^49, 50^ SVZ-born cells can also migrate outside of the RMS to repair damage in other regions of the brain.^51^ Disruption of these processes causes impairment in differentiation between olfactory stimuli and reduces overall brain plasticity.^49–51^

Chronic exposure to AgNPs was found to disrupt cognitive and behavioral function in mice.^26^ However, the long-term physiological impact of bioaccumulated AgNPs on cellular and synaptic mechanisms are less clearly defined. Here we demonstrate that subcellular nano-bio interactions could contribute to neurodegenerative mechanisms resulting from chronic environmental exposure to AgNPs and their bioaccumulation and persistence in brain tissue.

## RESULTS AND DISCUSSION

This study investigated whether oral AgNP exposure induces cellular mechanisms associated with neurodegeneration in the brains of young adult rats. Utilizing IHC and confocal microscopy, we found a significant increase in ptau and SOD1 localization to the nuclei of DG cells in hippocampus following exposure to acute and bioaccumulated AgNPs. In contrast, the overall proportion of phosphorylated tau and total SOD1 expression in hippocampus as measured by immunoblot were unchanged, implying specific nano-bio interaction mechanism in the cells of the DG. In addition, we describe disruptions in adult neurogenesis in the SVZ-RMS-OB niche, suggesting dysregulation of neural progenitor proliferation in response to acute and bioaccumulated AgNP exposure. These findings suggest that chronic exposure to environmental AgNPs, and their bioaccumulation and persistence in the brain, could be a risk factor for neurodegeneration and could be used to inform decisions on how AgNPs are incorporated into products, used in manufacturing and agriculture, and in waste management of products and water containing AgNPs.

### Characterization and validation of AgNPs

The AgNPs used in this study were commercially sourced. Emulsions were prepared fresh daily from dry powder for oral dosing of animals. Scanning electron microscopy (SEM) confirmed that individual AgNPs were spherical **(Fig. 1A)** with an average measured diameter of 40 nm (**Fig. 1B**). Mean size of individual particles as measured by dynamic light scattering was larger (∼198 nm), suggesting agglomeration that was also observed in the SEM images. Energy-dispersive X-ray spectroscopy (EDS) analysis confirmed the purity of the AgNP preparation with a strong silver spectral signature that was localized to individual nanoparticles (**Fig. 1C-G**).

**Figure 1:**
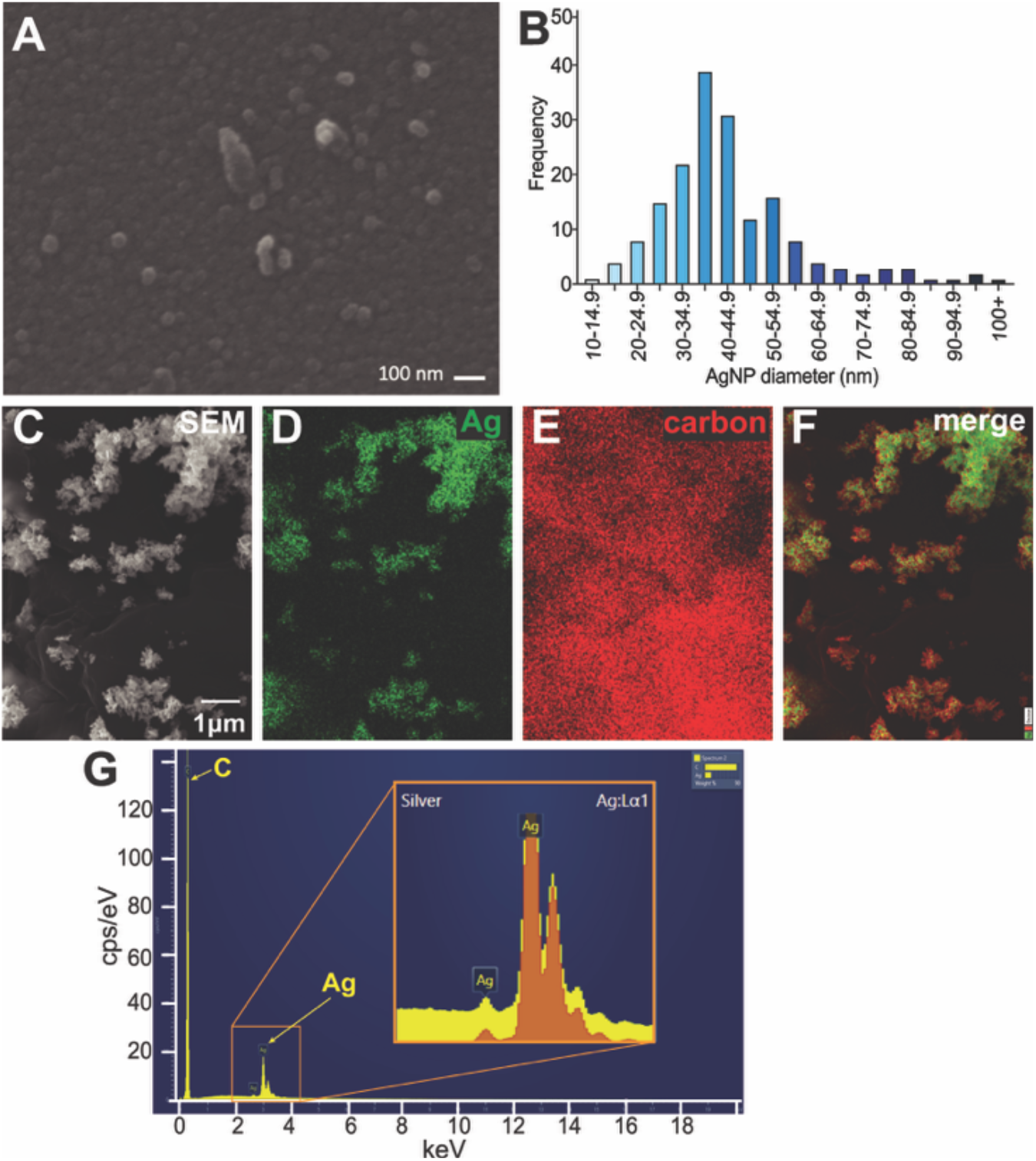
AgNPs fed to rats were confirmed to be pure Ag and have an average diameter of 40 nm. SEM of AgNP suspension dried to poly-lysine-coated coverslips (A). The diameter of 177 individual nanoparticles measured with ImageJ in SEM images confirm a frequency distribution centered around 35-40 nm (B). SEM of AgNP powder on a pure carbon stub (C) with elemental composition map of Ag (D), carbon (E), and merge (F). EDS spectrum of AgNP (yellow) superimposed with Ag reference spectrum (orange) in inset (G) confirms purity of AgNPs.

### Silver bioaccumulates and persists in rat brain after oral AgNP exposure

The popularity of AgNPs is rising in medicine as an alternative for combatting antibiotic-resistant bacteria^52^ and as a drug-delivery vehicle.^53^ They are also incorporated in a wide variety of consumer products including cosmetics, plastics like toothbrushes and toys, and textiles,^54^ yet the health consequences of these applications have yet to be fully understood. After oral AgNP exposure and brain collection, the cerebellum from each rat was analyzed to quantify elemental silver content using inductively coupled plasma mass spectrometry (ICP-MS). Ag accumulated in a dose-dependent manner in tissue collected the day after oral AgNP exposure (**Fig 2A, acute**). The mean Ag content after 9 weekdays of 25 mg/kg/day (acute) was 31.1 ng Ag/g tissue weight, and this increased to 61.2 ng Ag/g tissue weight in animals exposed to 100 mg/kg/day treatment. This translates to 3.1 and 6.8 ng Ag/g tissue weight/day for the low and high dose, respectively, suggesting that uptake may be nonlinear or saturated at higher exposure levels. In brain tissue collected from rats given the same nine-day AgNP exposure followed by a month without exposure (**Fig. 2A, bioaccumulated**), the Ag content was comparable to brains from acute treatment rats, with mean total Ag levels of 23.2 and 53.0 ng/g tissue weight after 25 and 100 mg/kg/day AgNP exposure, respectively. These data indicate persistence of bioaccumulated Ag in the brain for at least a month with no significant clearance. For some animals, Ag bioaccumulation in cerebellum was compared to that in a cerebral hemisphere **(Fig 2B),** confirming that Ag levels in cerebellum were representative of those in other brain regions.

**Figure 2.**
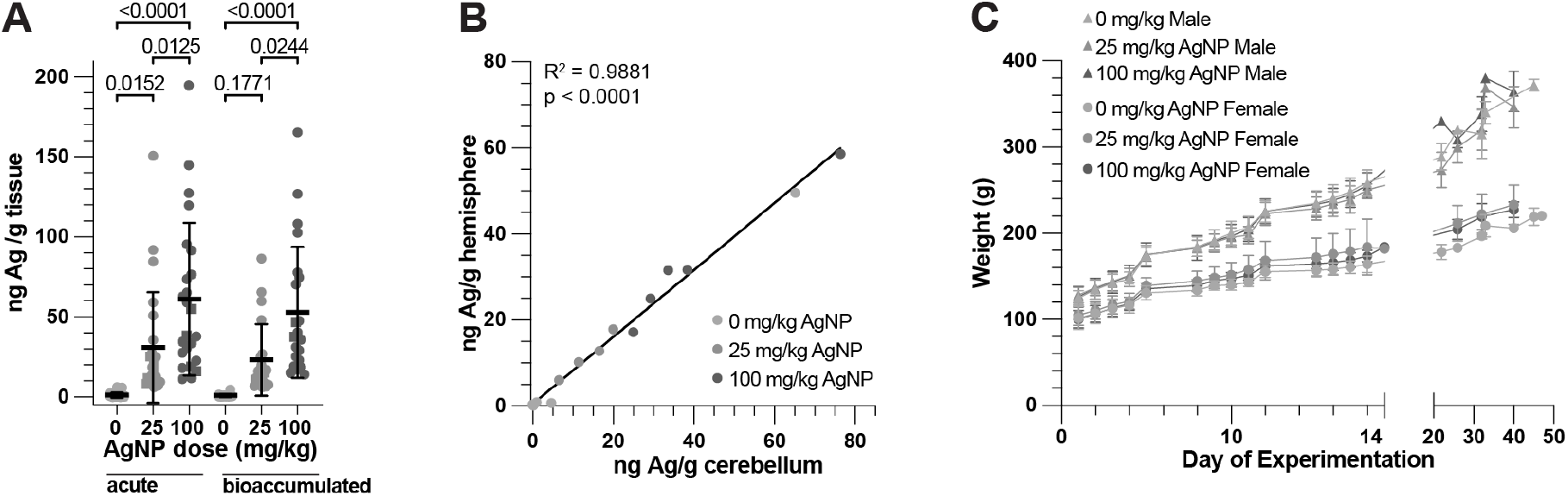
AgNP exposure accumulates and persists in the brain in a dose-dependent manner but does not to alter body weight. Cerebellum from rat brains collected immediately after AgNP treatment (acute) contain dose-dependent levels of Ag. These Ag levels were maintained in brains collected from rats kept for a month without additional exposure (bioaccumulation). N=21-24 animals per dose. One-way ANOVA followed by Tukey’s multiple comparisons. P values <0.05 are shown. (A). Silver concentration in a cerebral hemisphere is equal to that in the cerebellum for individual animals. N=5 animals for each AgNP dose. Simple linear regression. (B). Rat weight was unaffected by AgNP exposure throughout the experimental period N=8 cohorts. (C).

Our results are comparable to prior findings with similar treatments in rodents.^5, 21, 27–29, 55^ For example, a 28-day exposure to 10 nm or 25 nm AgNPs at 100 mg/kg/day resulted in a total accumulation of about 250 ng/g rat brain tissue, or about 9 ng/g tissue each day.^28^ Our study utilized 40 nm AgNPs over a shorter exposure period, which may account for our lower observed levels of bioaccumulation, as smaller particles may cross biological barriers more effectively.^35, 56^ It is also possible that bioaccumulation is not linear such that longer exposure results in a higher level of AgNPs incorporation by the measured endpoint. Indeed, AgNPs incorporated in brain tissue can serve as deposition centers, leading to nanoparticle growth during subsequent high Ag levels.^22^ Finally, we suspended the AgNPs in a sucrose solution for voluntary consumption instead of administration by gavage.^28^ It is possible that the sucrose vehicle altered uptake from the gut, perhaps by modifying the protein corona of the AgNPs, thereby modulating their interactions with the mucosal lining.^57^

Rat weight was unaffected by two weeks of AgNP exposure or during a month of exposure to bioaccumulated Ag (**Fig. 2C**), and overt behavioral changes were not observed, implying that AgNP exposure did not adversely affect overall animal health. Silver, in any form, is currently considered only as a secondary water quality standard by the environmental protection agency.^58^ Thus, the risk of accumulation and persistence of Ag in brain is a unique and growing challenge associated with the increased use of engineered AgNP in manufacturing and the release of these particles into the environment. Consumer goods can leach up to 18.5 mg/kg AgNPs,^10^ and human exposure to AgNPs has been estimated at 20-80 µg/day.^55^ Like others,^21, 28, 59^ we found that Ag persists in a dose-dependent manner over time despite no additional AgNP exposure, demonstrating bioaccumulation and persistence for at least one month. Notably, the bioaccumulation and persistence of Ag in the brain observed after exposure to engineered AgNPs did not occur when animals were exposed to ionic Ag.^60, 61^ Given the tendency of engineered AgNPs to enter and remain in brain tissue, it is important to understand how AgNPs, which could bioaccumulate and persist during chronic environmental exposure over years or decades, interact with neuronal physiology.

Ag released from nanoparticles can induce production of ROS and interrupt intracellular O_2_ reduction, causing cellular distress and intermittent apoptosis.^5^ Cytotoxicity of AgNPs depends on concentration and exposure period.^62^ Thus, exposure to low, noncytotoxic levels of AgNPs could still result in accumulation to concentrations that alter cellular physiology, especially if exposure is chronic. Additionally, short-term, high-concentration exposure models tend to underestimate true lifetime AgNP accumulation in organisms exposed to chronic, low-level environmental AgNPs.^15^ This implies that AgNP-resultant harm in short-term studies could be less adverse than that of realistic lifetime exposure. It has been argued that human environmental exposure is too low to cause damage in organs such as liver.^10^ However, the nature of brain structure and function is such that other regions may not readily be able to compensate if specific regions or synapses are impaired by local AgNP incorporation. Here, we report changes in brain suggesting the activation of neurodegenerative processes in animals exposed to AgNPs at a level that does not induce overt signs of toxicity.

### Acute AgNP exposure induces abnormal phosphorylation and nuclear translocation of tau and increases nuclear SOD1 expression in DG

AgNP exposure induces oxidative stress in multiple systems, including brain,^36^ that can induce transcription and nuclear translocation of SOD1 to protect chromatin from damage.^44^ Oxidative stress has also been linked to neurodegeneration, including the hyperphosphorylation of tau and formation of tau tangles.^63^ Although tau is classically recognized for its physiological role in axonal microtubule stabilization,^41^ increasing evidence suggests that abnormally hyperphosphorylated tau localizes to the nucleus, where it may disrupt chromatin organization, DNA protection, and regulation of cellular stress responses.^64^ The DG has high metabolic demand and remains one of the few brain regions that retains adult neurogenic capacity,^65^ characteristics that may increase vulnerability to AgNP-induced cellular stress. Therefore, we used IHC to measure the nuclear localization of ptau and SOD1 in the dentate gyrus of rats exposed to AgNPs. Brains collected one day after the 9-day oral AgNPs exposure paradigm, representing acute exposure, were stained to visualize ptau (**Fig. 3A-C**), SOD1 (**Fig. 3D-F**), and nuclei (**Fig. 3G-I**). Representative merged images (**Fig. 3J-L**) show a dose-dependent increase in ptau and SOD1 localization to DG nuclei. The normalized area of ptau label and intensity of SOD1 label were quantified in the granular layer of the DG in the dorsal hippocampus. Nuclear ptau was significantly increased in both the low and high exposure groups compared to control, with the highest elevation observed in animals exposed to 25 mg/kg/day (**Fig. 3M**). Conversely, nuclear SOD1 levels increased in a dose-dependent manner (**Fig. 3N).** These findings indicate that AgNP exposure disrupts tau phosphorylation homeostasis within a hippocampal subregion known to be particularly sensitive to oxidative and metabolic stress.^66^ Moreover, nuclear ptau area was lower in the 100 mg/kg/day AgNPs dose group compared to the 25 mg/kg/day exposure group, yet nuclear SOD1 was significantly elevated only in animals receiving the higher AgNP dose. These data indicate that the hippocampus may compensate to counteract tau hyperphosphorylation by increasing antioxidant pathway responses to AgNPs in a dose-dependent manner.

**Figure 3.**
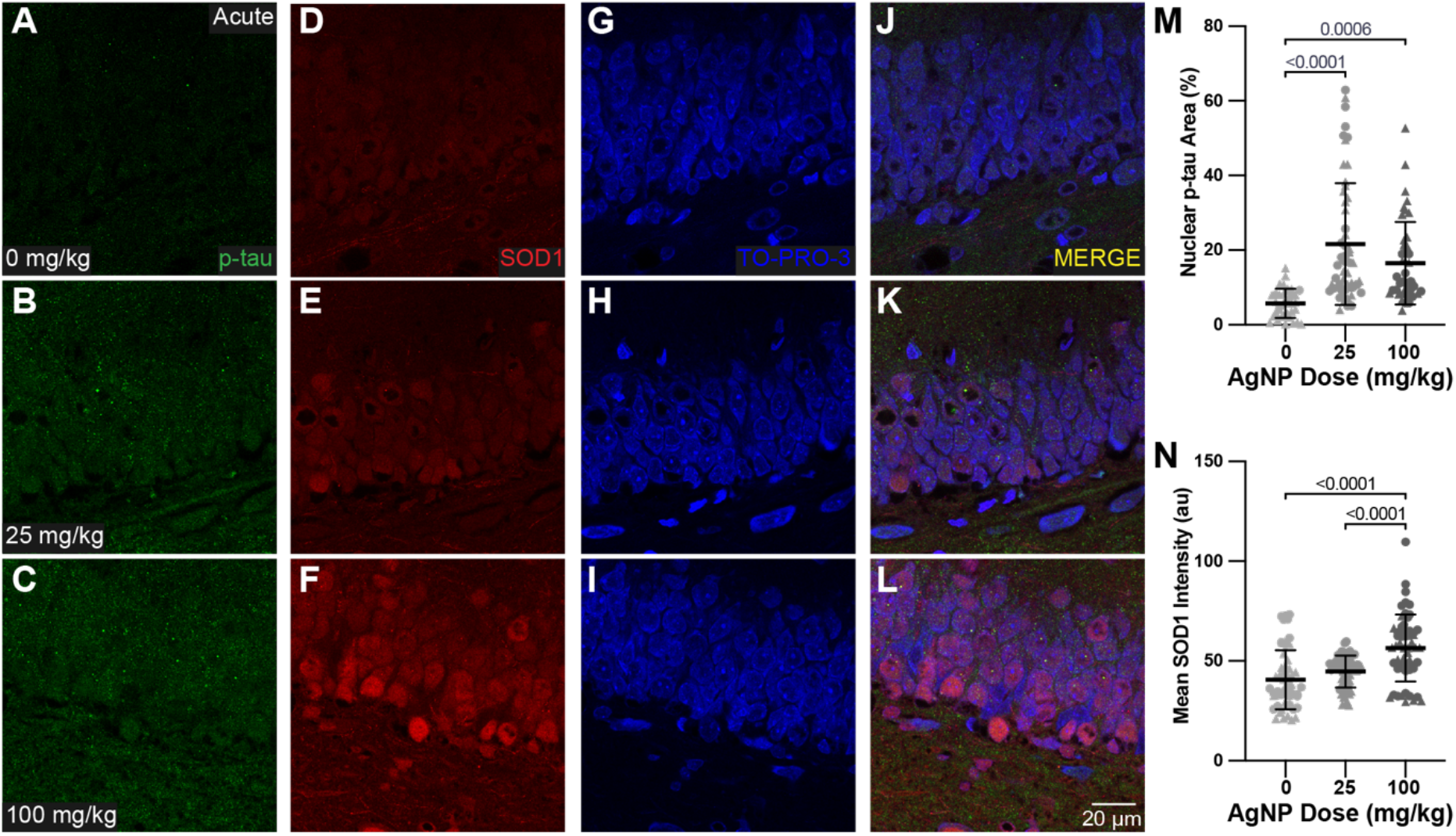
Nuclear localization of phosphorylated tau protein and increased SOD1 intensity in the granular layer of DG after acute AgNP exposure. Representative images of DG following nine consecutive weekdays of AgNP exposure to 0 (top), 25 (middle) or 100 (bottom) mg/kg/day (A-L). IHC was used to label ptau (green; A-C) and SOD1 (red; D-F) with TO-PRO nuclear counterstain (blue; G-H). Merged images (J-L) allow quantification of ptau area normalized to total nuclear area (M) and SOD1 intensity (N) in nuclei. Each experimental group contained N=7 animals with four females (circles) and three males (triangles). Data are represented as mean ±. S.D. One-way ANOVA followed by Tukey’s multiple comparisons. P values < 0.05 are shown. Scale bar 20μm.

The observed increase in nuclear ptau within the DG following AgNPs exposure may reflect altered nuclear homeostasis. Aberrant tau phosphorylation has been associated with impaired tau function,^67^ altered intracellular localization, and disruption of signaling pathways involved in cellular stress response. Nuclear tau interacts with DNA, chromatin, and other critical nuclear components, ^64^ thus accumulation of ptau in the nucleus could lead to transcriptional changes and alterations in neuronal activity or cellular stress resilience. The localization of ptau to nuclei within the granular layer of DG is of concern, given that the DG serves as a critical bridge for information entering the hippocampus and is involved in encoding and discrimination of contextual and spatial information.^68, 69^ When taken together, the nuclear accumulation of ptau with the increase in SOD1 expression may indicate an enhanced antioxidant response in DG cells associated with altered redox homeostasis. Oxidative balance is crucial to maintaining neuronal signaling and synaptic plasticity. Persistent alterations in redox homeostasis or nuclear ptau accumulation induced by chronic exposure to AgNPs could therefore change the ability of cells within the DG to maintain transcriptional responses, synaptic plasticity, and neuronal adaption^70^ and may lead to disruption of the mechanisms underlying learning and memory. Indeed, exposure to daily AgNPs over four months was correlated with poor long-term and contextual memory outcomes in mice.^26^

### Abnormal phosphorylation and nuclear translocation of tau and elevated nuclear SOD1 expression persist following exposure to bioaccumulated AgNP

Because we found that Ag persists in the brain for at least one month following exposure with no significant clearance, we tested the effect of exposure to bioaccumulated AgNPs on ptau (**Fig. 4A-C**) and SOD1 (**Fig. 4D-F**) in the nuclei (**Fig. 4G-I**) of DG cells in brains collected from rats one month after the acute exposure was complete. Merging of images (**Fig. 4J-K**) allowed measurement of nuclear ptau and SOD1. The area of nuclear ptau remained significantly elevated following exposure to bioaccumulated AgNPs at both the 25 and 100 mg/kg/day dose compared to animals that received sucrose vehicle only (**Fig. 4M**). In parallel with increased ptau, SOD1 expression was also significantly elevated following exposure to bioaccumulated AgNPs, with the highest elevation in nuclear SOD1 found in brains of rats that received 25mg/kg/day (**Fig. 4N**). In brains of rats that received 100mg/kg/day, SOD1 expression was slightly reduced compared to the 25mg/kg/day group but remained significantly elevated compared to control (**Fig. 4N**). These data suggest that activation of cellular stress responses are maintained in animals exposed to bioaccumulated AgNP for a month after the initial exposure.

**Figure 4.**
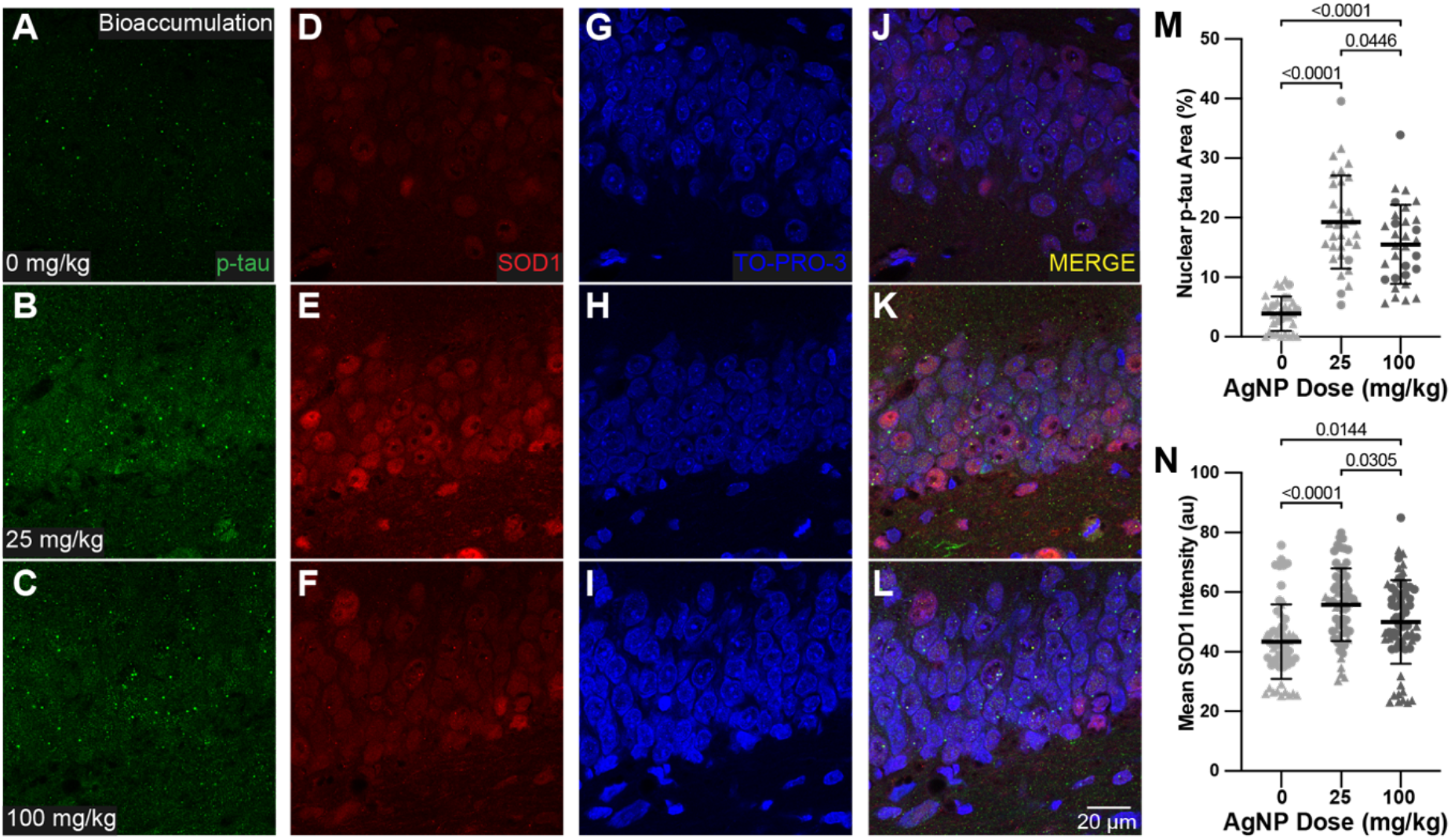
Nuclear localization of ptau and SOD1 expression in the nuclear layer of DG is further increased after exposure to bioaccumulated AgNPs. Representative images of DG following nine consecutive weekdays of AgNP exposure to 0 (top), 25 (middle) or 100 (bottom) mg/kg/day followed by one month of no additional exposure. IHC was used to label ptau (green; A-C) and SOD1 (red; D-F) with TO-PRO nuclear counterstain (blue; G-H). Merged images (J-L) allow quantification of ptau area (M) and SOD1 intensity (N) in nuclei. ptau area was normalized to the total nuclear area. Each experimental group contained N=7 animals with four females (circles) and three males (triangles). Data are represented as mean ± S.D. One-way ANOVA followed by Tukey’s multiple comparisons. p values < 0.05 are shown. Scale bar 20μm.

SOD1 is a key antioxidant enzyme involved in the detoxification of ROS, and its upregulation is consistent with activation of an oxidative stress response. Oxidative stress has been implicated in the regulation of tau phosphorylation through modulation of redox-sensitive kinase and phosphatase pathways, including GSK3β, CDK5, and PP2A signaling.^39^ Thus, the concurrent elevation of SOD1 and ptau observed in the present study suggests that oxidative stress may contribute to AgNP-induced dysregulation of tau phosphorylation. Phosphorylation at Ser396 of tau has been identified among the earliest events in neurodegenerative diseases involving tau pathology.^67^ Increased phosphorylation at Ser396 may reflect activation of stress-responsive GSK3β and CDK5 signaling cascades which are known to regulate tau pathology.^71^ Importantly, the persistence of elevated ptau after one month of exposure only to bioaccumulated AgNPs suggests that antioxidant SOD1 elevation and other potential compensatory mechanisms to prevent sustained dysregulation of cellular homeostasis are not sufficient to counteract the disruptions induced by bioaccumulated AgNPs persisting in the brain.

Collectively, these findings demonstrate that exposure to AgNPs induces sustained elevations in nuclear ptau and antioxidant responses within the DG. The persistence of these molecular alterations following exposure to bioaccumulated AgNPs highlights the potential for chronic exposure to environmental AgNPs to produce lasting effects on hippocampal cellular homeostasis, even after removal of the exposure source. Although the present study did not assess downstream functional consequences, the persistence of nuclear ptau and SOD1 elevation raises the possibility that chronic AgNP exposure, in combination with bioaccumulation and persistence, could induce prolonged alterations in hippocampal molecular signaling.

Toxicologically, this distinction is critical, as removal of AgNP exposure did not result in clearance of the AgNPs themselves, or a return to baseline cellular function, for as long as one month after ingestion. Persistent changes in oxidative stress and tau regulation pathways in response to chronic environmental exposure to AgNPs may therefore increase the risk of neurodegeneration, especially when combined with additional environmental,^72^ physiological,^73^ or genetic^74^ risk factors.

### Total ptau and SOD1 in hippocampus are unchanged by acute AgNP exposure but increased after exposure to bioaccumulated AgNPs

To assess more global changes in tau hyperphosphorylation associated with exposure to AgNPs, immunoblot was used to quantify the proportion of ptau to total tau protein in the dorsal hippocampus (**Fig. 5**). In contrast to the IHC findings reported above, immunoblot demonstrated no change in tau phosphorylation in hippocampus after acute AgNP exposure (**Fig. 5A&E**). However, after exposure to bioaccumulated AgNP, the total hippocampal proportion of ptau was increased (**Fig. 5C&G**), suggesting that bioaccumulated AgNP could lead to broader dysregulation of tau homeostasis. Overall, AgNP-induced effects on the proportion of ptau were not as substantial when measured by immunoblot compared to IHC, suggesting that AgNP interactions with mechanisms regulating tau phosphorylation and nuclear translocation may be specific to DG cells, and not necessarily global across the region. These findings raise the possibility that nuclear ptau accumulation represents a localized response to AgNP exposure that may lead to specific neurodegenerative risks. Immunoblot for total SOD1 expression showed variable SOD1 levels in hippocampus from rat brains collected after acute AgNP exposure (**Fig. 5B&F**). Consistent with IHC results, there was a dose-dependent increase in the proportion of ptau protein in hippocampus from rats exposed to bioaccumulated AgNPs (**Fig. 5C&G**). This discrepancy between acute or bioaccumulated AgNPs tissue may reflect a temporally dynamic cellular response to AgNP exposure where acute exposure may initially promote redistribution or compartmentalization of ptau, including nuclear translocation within the DG, without adding to total hippocampal ptau abundance. As in hippocampus from rats exposed to acute AgNP, total SOD1 levels measured by immunoblot were variable and relatively unchanged compared to control in animals exposed to bioaccumulated AgNPs (**Fig. 5D&H**). As with ptau, these findings suggest that AgNP-induced activation of antioxidant pathways is specific and localized instead of global across the entire brain region. Thus, the spatial resolution of IHC may more accurately represent changes in nuclear ptau and SOD1 observed after exposure to AgNPs.

**Figure 5.**
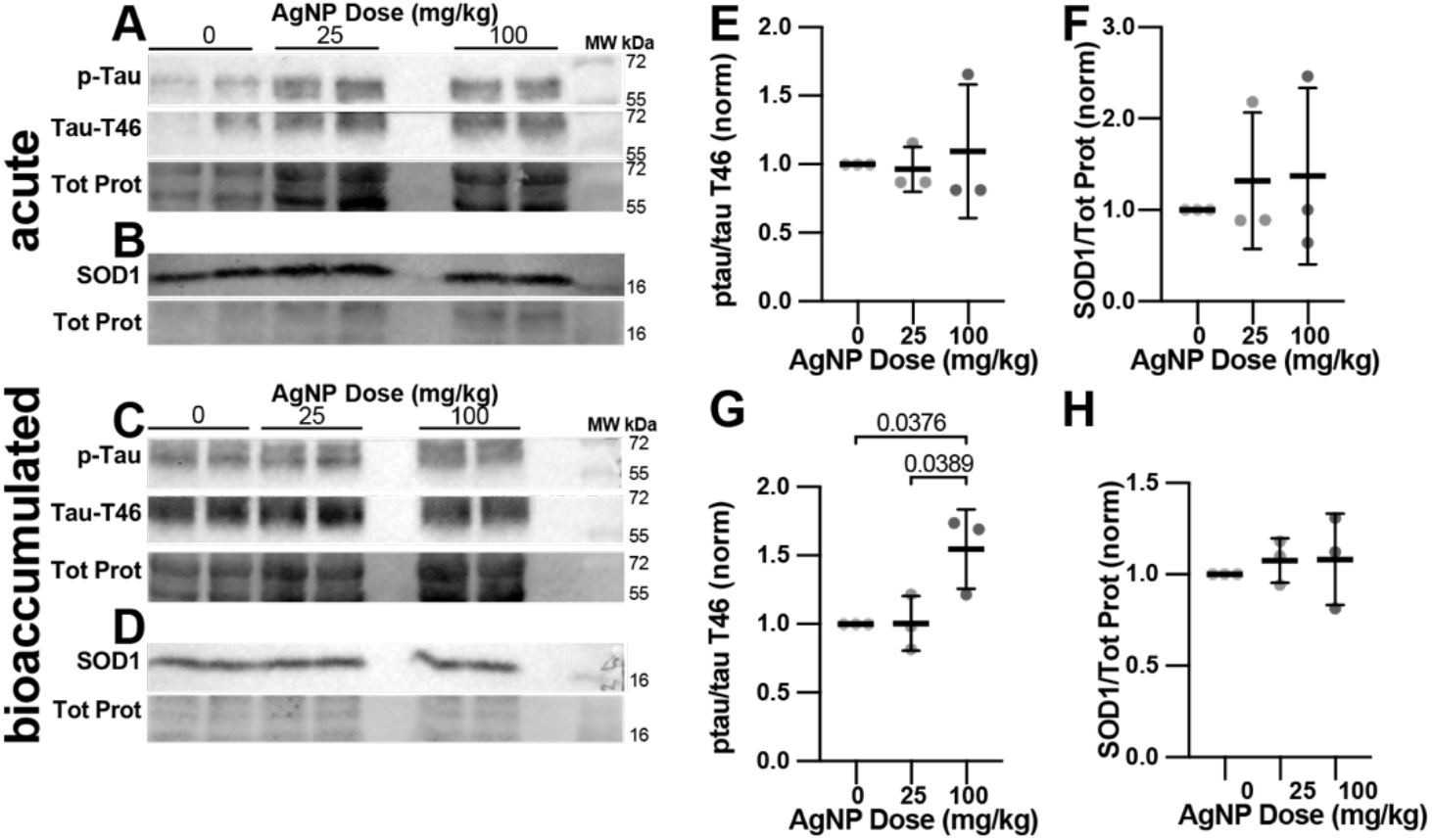
Proportion of ptau increases across global hippocampus after exposure to bioaccumulated AgNPs. Representative images of immunoblots conducted on hippocampal tissue after acute AgNP treatment probed for ptau, pan tau (tau T46), and SOD1, with total protein (A&B). Representative images of immunoblots conducted on hippocampal tissue after exposure to bioaccumulated AgNPs probed for ptau, pan tau, and SOD1, with total protein (C&D). The proportion of ptau compared to tau T46 (pan) (E) and expression of SOD1, normalized to total protein, (F) remained unchanged following acute AgNP exposure. The proportion of ptau was significantly increased after exposure to 100 mg/kg bioaccumulated AgNP (G), but SOD1 expression remained unchanged (H). Data are represented as mean ± SD. One-way ANOVA followed by Tukey’s post hoc. N=3 animals for each condition.

### Acute AgNP exposure induces disruption of neurogenesis in the SVZ-RMS-OB system

We previously found that exposure to AgNP disrupted f-actin cytoskeleton, induced neurite collapse, and altered specific intracellular signaling pathways in cultured differentiating adult-derived neural stem cells and neural cell lines.^30–33^ Therefore, we sought to investigate the effect of oral AgNP exposure on adult neurogenesis in the SVZ-RMS-OB niche in the young adult rat brain. Neuroblasts are born in the SVZ and migrate along the RMS, continuing to proliferate, until they are incorporated in the olfactory bulb where they complete their maturation and integrate into existing circuits.^49^ All of these processes require active cytoskeletal remodeling and control.

The thiamine analog, 5-ethynyl-2’-deoxyuridine (EdU), was used to mark the nuclei of new cells in the brain, and the EdU was labeled with a red fluorescent label using click chemistry for detection (**Fig. 6A**). EdU incorporation was measured in three regions along the RMS: at the SVZ (**Fig. 6B**), the rostral bend (**Fig. 6C**), and the end of the RMS at the OB (**Fig. 6D**). Neurogenesis within each region was quantified as the area of EdU label within the RMS as delineated by all nuclei visualized with DAPI. Because EdU labels only nuclei, which are of uniform size, area was used as a proxy for the number of new cells in each image. We found high variability in the degree of EdU incorporation across all treatment conditions and all regions of the RMS. We used a single dose of EdU to label dividing cells in an effort to limit our investigation to cells that divided only in the 2-day window after EdU injection. However, this did not allow us to differentiate fully between cells that were born earlier and still dividing along the RMS and those that were born in the SVZ during the EdU exposure; these cells are able to travel about halfway along the RMS in 2 days.^48^ This means that slight variation in either proliferation or migration mechanisms could result in great data variability overall. We found no change in the number of new cells at the SVZ in acute AgNP treatment animals (**Fig. 6E1-3 & K**), suggesting that the rate of neurogenesis and migration from the SVZ is not impacted immediately following AgNP exposure. Similarly, the number of new cells in the rostral bend of the RMS was unchanged in animals with acute AgNP exposure (**Fig. 6F1-3 & L**). Newly born cells from the SVZ are expected to migrate about halfway along the RMS, to the rostral bend region, in the two days after EdU injection,^48^ suggesting that the labeled population in this region represents a mixture of cells that migrated from the SVZ after the label was injected and those that may have divided locally. At the OB end of the RMS, a dose-dependent increase in new cells was observed (**Fig. 6G1-3 & M**), with rats exposed to 25 mg/kg/day having significantly more EdU^+^ cells in the RMS at the OB compared to control. The increase in new cells in response to 100 mg/kg/day also reached significance when outlier data points were removed. Given the rate of migration along the RMS,^48^ labeled cells in the OB are likely the product of cell division within the RMS more than migration of labeled progenitors from the SVZ. The increased numbers of new cells at the OB suggest that acute AgNP exposure may promote cell division of new neural stem cells as they migrate along the RMS. Such an increase in neurogenesis has been observed in response to exercise^75^ and physiological insults including epilepsy^76^ and traumatic brain injury.^77^ Thus, acute AgNP exposure may induce compensatory protective mechanisms leading to increased neurogenesis in the SVZ-RMS-OB niche. In animals exposed to bioaccumulated AgNPs for a month after acute exposure, numbers of new cells were no different from control at the SVZ (**Fig. 6H1-3 & N**). As in acute exposure rats, this indicates that neurogenesis and migration in the SVZ are unchanged by exposure to bioaccumulated AgNPs. However, in the rostral bend (**Fig. 6I1-3 & O**) and at the OB (**Fig. 6J1-3 & P**), the trend was opposite to what was observed after acute exposure, with the RMS containing a reduced number of new cells after exposure to bioaccumulated AgNPs compared to control, suggesting that compensatory mechanisms activated by acute exposure may not persist during chronic exposure which could lead to a neurogenesis deficit. The data are highly variable, suggesting that local cell division in the RMS may be inhibited by exposure to bioaccumulated AgNP. Alternatively, newly proliferated cells in the RMS could be leaving and migrating to other regions of the brain to counteract AgNP-induced damage,^51^ leading to an apparent loss in RMS neurogenesis. We may also be observing a shifting balance between AgNP-induced damage and compensatory mechanisms, such as increased antioxidant activity. Apoptosis is active in ‘pruning’ newly generated cells in the SVZ-RMS-OB system,^78^ and AgNP exposure could inappropriately enhance this mechanism. In traumatic brain injury, neurogenesis can significantly increase,^77^ therefore, depending on the efficacy of compensatory mechanisms, this could account for the dichotomy in neurogenesis levels between the acute and bioaccumulated exposure. This may also account for the apparent increase in proliferation at the end of the RMS following acute exposure. Many physiological factors drive the level of neurogenesis in the SVZ-RMS-OB system, and this niche is highly plastic in young adult animals, therefore high variability and complex effects are not unexpected. More targeted *in vivo* labeling approaches or cell culture may provide more insight on the interaction of acute and bioaccumulated AgNPs with neural progenitors in the brain.

**Figure 6.**
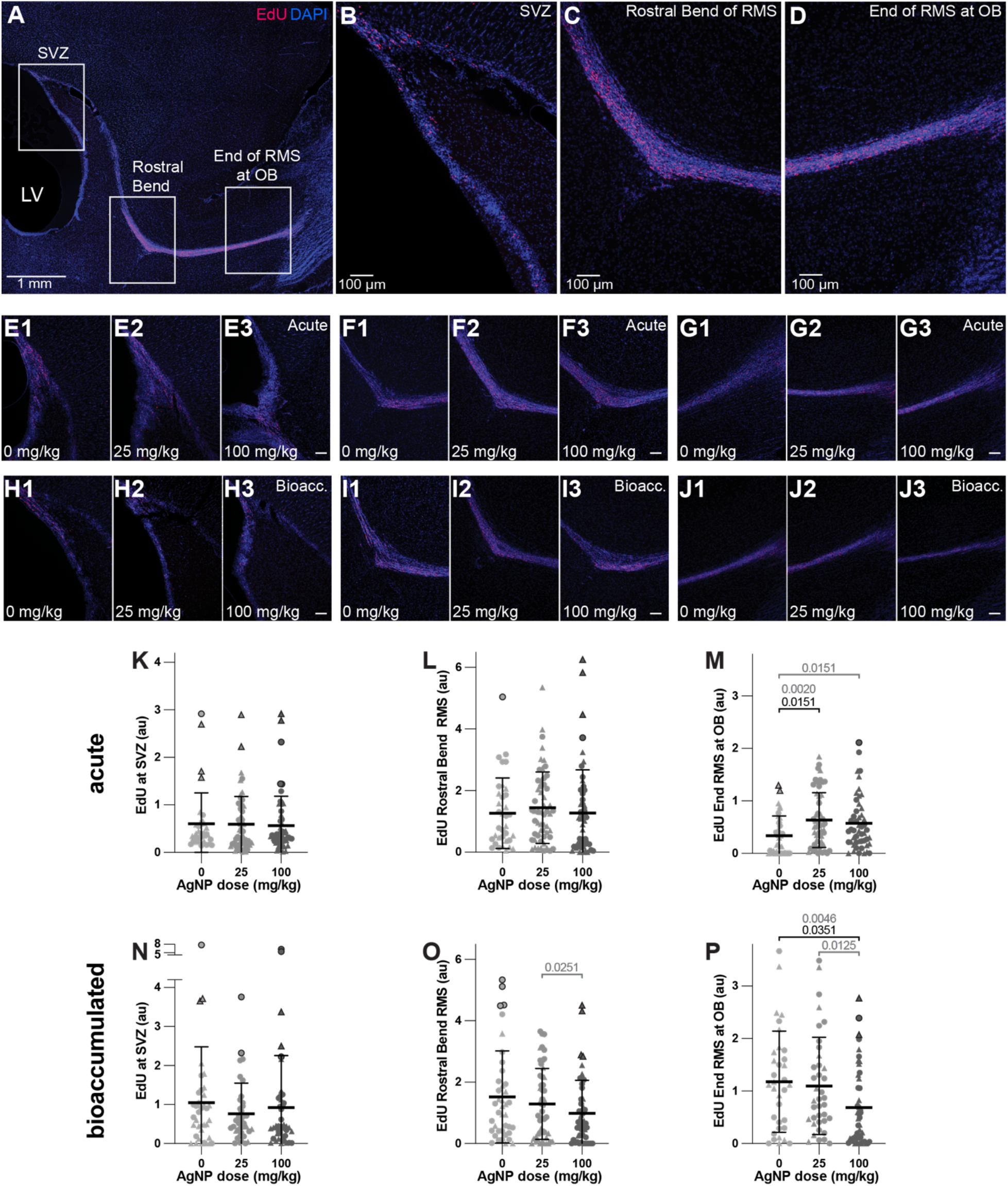
Exposure to AgNPs only slightly alter neurogenesis along the RMS but changes the number of newborn cells reaching the OB. Nuclei of newly born cells in the SVZ-RMS-OB system (A) were labeled with EdU (red), and all nuclei were visualized with DAPI (blue). Images were collected at the SVZ (B), at the rostral bend (C), and at the OB end (D). Representative images are shown for the SVZ (E1-3), rostral bend (F1-3), and the OB end (G1-3) in brains collected immediately after AgNP treatment (acute) and the SVZ (H1-3), rostral bend (I1-3), and the OB end (J1-3) in brains collected from animals that were exposed to bioaccumulated AgNPs for one month after treatment. The area of EdU label was quantified after acute AgNP treatment from the SVZ (K), at the rostral bend (L), and at the OB (M). Corresponding analyses were performed at the SVZ (N), rostral bend (O), and OB (P) in brains collected from animals that were exposed to bioaccumulated AgNPs. Up to five images were analyzed from each of N=7-11 animals including 3-5 females (circles) and 4-6 males (triangles). Outlier data points (dark outlines) were identified by ROUT with Q=1%. Data are represented as mean ± SD calculated with outliers included. One-way ANOVA followed by Tukey’s multiple comparisons. P values <0.05 are shown in black for full data and in grayscale for calculations with outliers removed. Scale bars 1mm in (A) and 100μm in (B-J).

## Conclusion

In summary, our data indicate that oral exposure to AgNPs induces mechanisms associated with cellular stress and neurodegeneration in a brain region important for learning and memory. Moreover, AgNPs persist in the brain a month after cessation of exposure and neurodegenerative markers remain elevated, suggesting that compensatory mechanisms are not sufficient to counteract local cellular disruptions resulting from bioaccumulated persisting AgNPs. These disruptions appear to be specific and localized as global immunoblot measurements of neurodegeneration and oxidative stress were altered much less severely. Similarly, only minor changes were detected in the complex SVZ-RMS-OB adult neurogenesis niche. Taken together, chronic exposure to low-level environmental AgNPs may induce bioaccumulation and persistence of these particles in the brain, which may lead to increases in risk for neurodegenerative disorders.

## METHODS

### Nanoparticle Characterization and Preparation, Dosage, Exposure, and Tissue Collection

AgNPs (40 nm) were purchased as a dry powder (Nanostructured and Amorphous Materials, Inc., Katy, TX). For characterization, AgNPs were suspended in tetrahydrofuran at 1 mg/ml. They were sonicated for 15 min and then analyzed by dynamic light scattering (DLS; Brookhaven) or dried on coverslips and atomic carbon coated for scanning electron microscopy (SEM; JEOL). Dry powder was directly applied to a pure carbon mount (SPI supplies) and energy-dispersive X-ray spectroscopy (EDS) was performed using the Ultim Max attachment from Oxford Instruments. EDS Spectra were collected, and elemental peaks identified using AztecLive Spectrum acquisition and X-Ray smart mapping software. Spectra were collected with an emission of 20.0 Kev at a working distance of 10.0 mm.

For oral dosing of animals, AgNP solutions were prepared fresh from dry powder each day. Water was added to AgNPs to make a 100 mg/ml solution. A probe sonicator was used to solubilize the AgNPs with three bursts of 20 KHz at an amplitude of 144 µm for three min separated by three min rests on ice. The resulting solution was diluted in half with 20% sucrose in water to obtain 50 mg/ml AgNP in 10% sucrose for the high treatment. An aliquot of this 50 mg/ml solution was further diluted to 12.5 mg/ml with 10% sucrose to make the low AgNP treatment. Some agglomeration in the solution (as detected by DLS and SEM) is not a concern because individual particles are expected to be emulsified as they pass through the digestive tract,^3^ allowing for absorption in the gut. Control animals received the 10% sucrose vehicle solution only. Animals received 0 mg/kg, 25 mg/kg, or 100 mg/kg, respectively, each day for nine consecutive weekdays. This approach was informed by Colman et al.,^15^ who showed that repeated and temporally distributed AgNP exposure can produce sustained tissue burdens and delayed biological effects distinct from acute exposure alone. Although administered doses were expressed in mg/kg body weight, the biologically relevant exposure metric for tissue-level analyses were Ag accumulation within the brain, quantified as ng/g tissue. Systemic dosing does not directly translate to equivalent tissue concentrations because nanoparticle biodistribution is influenced by absorption, circulation, BBB transport, clearance kinetics, and tissue retention.

A cohort consisted of six rats that were split into two sets of three experimental groups: control (vehicle only), low AgNP exposure (25 mg/kg/day), and high AgNP exposure (100 mg/kg/day). Eight cohorts, half male and female, were used for the ptau/SOD1 experiments and twelve cohorts, half male and female, were completed for neurogenesis analysis. Immunoblot analysis was performed on three cohorts (2 male, 1 female). Upon arrival, rats were given one week to acclimate. To train rats to consume the AgNP solution, they were given a 10% sucrose solution daily for five days. Once rats completed sucrose training, they received AgNPs emulsified in 10% sucrose for nine consecutive weekdays. Solutions were consumed by the animals voluntarily. One set of brains was collected one day after AgNP exposure (acute), and the second set was collected one month after AgNP exposure to test the effects of bioaccumulated AgNPs (bioaccumulation).

For neurogenesis analysis, rats were lightly anesthetized with 5% isoflurane in oxygen via a nose cone and vaporizer (Kent Scientific, Torrington, CT) two days prior to brain collection. An intraperitoneal injection of 10 mg/mL sterile 5-ethynyl-2’-deoxyuridine (EdU; Carbosynth, UK) in 0.9% NaCl was administered at 50 mg/kg. Once injected, EdU incorporates into the DNA of newly proliferated cells.

For tissue collection for immunohistochemistry and click reaction staining, rats were deeply anesthetized with 5% isoflurane and perfused with phosphate-buffered saline (PBS1; 2.7 mM Na_2_HPO_4_, 9.2 mM NaH_2_PO_4_, 137 mM NaCl, 27 mM KCl, pH 7.4) followed by 4% paraformaldehyde (PFA) in PBS1 for fixation. Each brain was post-fixed in 4% PFA for 24 hr at 4°C then switched to 30% sucrose in PBS1 at 4°C. After the tissue was equilibrated, brains were cut into left and right hemispheres and frozen in 30% sucrose in PBS1/OCT (2:1) at -70°C. The hemispheres were sectioned sagitally with a cryostat at 30μm, lateral to medial. Free-floating sections were stored in cryoprotectant (30% ethylene glycol, 30% glycerol, in PBS2 (20 mM NaH_2_PO_4_, 80 mM Na_2_HPO_4_, 152 mM NaCl, pH 7.4)) at -20°C. For immunoblot, animals were deeply anesthetized with 5% isoflurane and decapitated with a guillotine. The brain was quickly collected, and the dorsal hippocampus was dissected and flash frozen in liquid nitrogen. Tissues were stored at -70°C. All work was carried out with the formal approval of the Marshall University Institutional Animal Care and Use Committee.

### Click Reaction Staining for EdU Incorporation and Immunohistochemistry

To quantify neurogenesis in the SVZ-RMS-OB system, all sections containing the rostral curve of the RMS from one hemisphere were processed through click chemistry to label nuclei with incorporated EdU. Sections were placed in mesh wells and washed in PBS2 4 times for 5 min on a rocker. To permeabilize and block the tissue, sections were transferred into a solution of 3% bovine serum albumin (BSA; Fisher) in 0.3% Triton-X/PBS2 (PBTx) once for 5 min and again for 30 min. The sections were then washed in PBS2 4 times for 5 min. To label EdU, the sections were placed for 30 min in an azide click reaction solution (0.1 M tris-buffered saline pH 7.6, 0.04 M CuSO_4_, 0.1 M sodium ascorbate) with a red Sulfo-Cyanine3 azide (Lumiprobe, Hunt Valley, MD) fluorophore kept in the dark. Tissues were kept in the dark for the remainder of the labeling process. After the click reaction, sections were transferred directly into a 3% BSA in PBS2 solution for 10 min. The sections were washed in PBS2 4 times for 5 min and then placed in a 1:1000 4’,6-diamidino-2-phenylindole (DAPI; Sigma) in PBS2 solution for 5 min to label nuclei.

For immunohistochemistry, brain sections were washed with PBS2 and blocked in 10% PBTx, 10% normal goat serum (NGS), and 1% donkey serum for one hour. Next, the tissue was moved into the primary antibody solution containing: 1:250 rabbit anti-ptau S396 (Invitrogen #44-752G) and 1:100 sheep anti-SOD1 (Millipore #574597) in block solution. After an overnight incubation with primary antibodies at 4°C, sections were washed 6 times in PBS2. Sections were then washed and incubated with secondary antibodies: goat anti-rabbit-Alexa 488, and donkey anti-sheep Alexa-568 at 1:200 each. On day three, all sections were washed with PBS2 and stained with 1:250 TO-PRO-3 (Invitrogen #R37113) in PBS2 to label nuclei. Free floating sections were then mounted and cured using Prolong Glass (Invitrogen).

### Immunoblot

Hippocampal tissue was homogenized in RIPA buffer (Thermo) using a BeadBug (Benchmark) and diluted with SDS Page loading buffer (Cell Signaling Technology) and aliquots were stored at -70°C. Samples were separated on a stain-free gel (BioRad TGX Stain-Free FastCast Acrylamide Kit, 10% #1610183). Gels were imaged for total protein and then transferred to a PVDF membrane (Thermo). Total protein was imaged again on the membranes to confirm transfer and allow normalization of chemiluminescent signals to total protein loaded. Membranes were blocked with 4% milk in tris-buffered saline with 0.1% tween-20 (TTBS), washed in TTBS and then probed with primary antibodies rabbit anti-p-Tau (1:1000; Invitrogen #44-752G) or sheep anti-SOD1 (1:100; Millipore #574597) in 5% BSA/TTBS solution and rocked at 4°C overnight. Membranes were washed with TTBS and incubated for one hour in secondary antibodies 1:1000 goat anti-rabbit HRP (1:1000) or, donkey anti-sheep HRP (1:1000) in 5% BSA/TTBS. Membranes were developed with ECL reagent (ThermoScientific #32209) and images acquired with a ChemiDoc system (BioRad). Membranes were then washed with TTBS and stripped with Restore buffer (Thermo). Total protein was imaged again, the membranes were re-blocked for an hour in 4% milk/TTBS, and then incubated with mouse-anti-TauT46 (1:1000; Invitrogen # 13-6400) and developed as described above.

### Imaging and Analysis

For ptau and SOD1, confocal Z-stacks of fifteen images each were acquired from two brain sections for every animal with a Leica Stellaris 8 microscope. Fiji (ImageJ; NIH, Bethesda, MD) was used to quantify ptau and SOD1. Z-stacks were split into five projections consisting of three images each to ensure no overlap of nuclei. We generated a macro in Fiji to automatically create a Z projection (**Fig. 7A**), split the channels, binarize and outline the nuclei to generate a region of interest (ROI) around the DG (**Fig. 7B-C**). Within this ROI, the macro then measured the area of ptau aggregates (**Fig. 7D**). SOD1 expression was analyzed using a similar workflow to delineate nuclei and measure the mean intensity of SOD1 label within that nuclear ROI.

**Figure 7.**
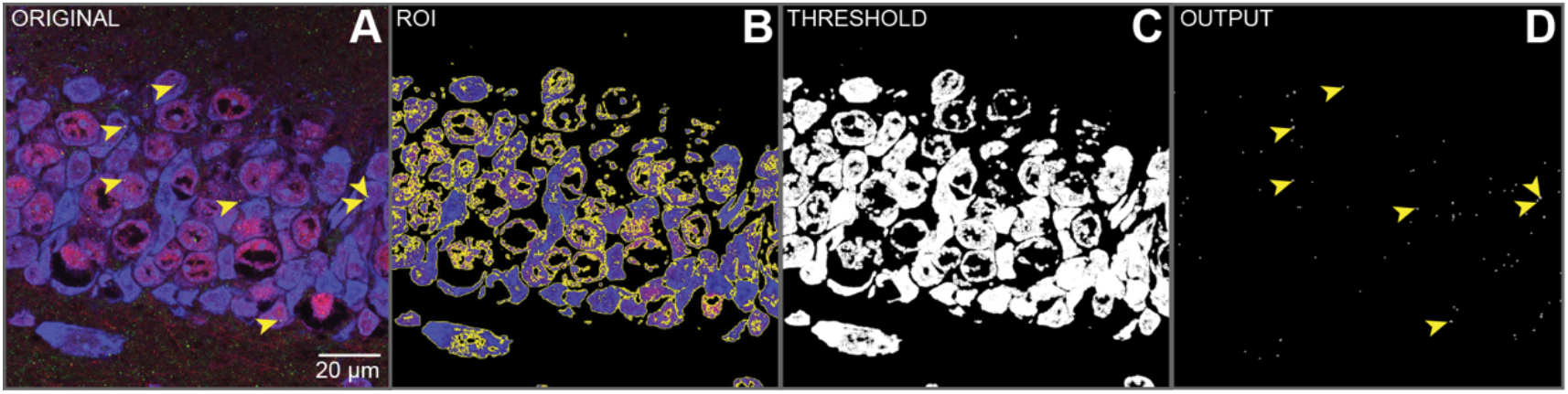
Analysis workflow for ptau analysis. Original image is opened in Fiji (A). The macro outlines the nuclear layer of dentate gyrus (B) and binarizes to clear outside to generate ROI (C). ROI is applied to green channel (ptau) and macro quantifies area of ROI that contains ptau (D).

EdU label was imaged with an inverted epifluorescent microscope (Leica, Germany) in Z-stacks spanning the full width of the RMS at three locations: the SVZ, the rostral bend, and the end of the RMS at the OB. Fiji was used to quantify EdU incorporation. An ROI was outlined around the RMS using the DAPI signal. Blue DAPI and red EdU labels were thresholded, and their respective areas were measured as above.

Immunoblot images were analyzed using BioRad ImageLab software. Samples from three cohorts were run in duplicate and intensity averaged. The proportion of ptau band intensity over pan-tau (T46) for each lane was calculated and SOD1 band intensity was normalized to stain-free total protein for that lane, the average for each sample calculated as a single data point.

Prism (Graphpad) was used to graph data and statistical analysis using one-way ANOVA followed by Tukey’s multiple comparisons. P values <0.05 are shown in black. For Figure 6, outliers were identified using the ROUT method with Q=1%. Outlier data points are shown with dark outlines and mean ± SD calculations on graphs include outlier data points. Statistical comparisons with outliers removed are shown in grayscale.

## AUTHOR INFORMATION

## Author Contributions

The manuscript was written through contributions of all authors. All authors have given approval to the final version of the manuscript.

## ACKNOWLEDGMENT

We thank A.M. Bora, E.S. Williams, and E.J. Peterson for aid in treating rats, tissue prep, and making graphs of rat weights and Dr. M-L. Risher, O. Coulter, T. Muck, and Dr. C. Risher for help with tissue preparation and processing protocols. E. Yablonsky helped collect some EdU images. Thank you to Dr. R. Quiñones of the Chemistry Department at Marshall University for DLS analysis. A. Lutton and J. Olesik at the Ohio State University Trace Element Research Laboratory performed ICP-MS analysis. SEM-EDS analysis was performed on an instrument supported by NSF #CHE1828358 at the Marshall University Molecular and Biological Imaging Center with training and help from D. Neff.

## ABBREVIATIONS

AgNPs: silver nanoparticles;
ptau: phosphorylated tau;
DG: dentate gyrus;
SOD1: superoxide dismutase 1;
OS: oxidative stress;
ROS: reactive oxygen species;

